# Plasticity at the DNA recognition site of the MeCP2 mCG-binding domain

**DOI:** 10.1101/581934

**Authors:** Ming Lei, Wolfram Tempel, Ke Liu, Jinrong Min

## Abstract

MeCP2 is an abundant protein, involved in transcriptional repression by binding to CG and non-CG methylated DNA. However, MeCP2 might also function as a transcription activator as MeCP2 is found bound to sparsely methylated promoters of actively expressed genes. Furthermore, Attachment Region Binding Protein (ARBP), the chicken ortholog of MeCP2, has been reported to bind to Matrix/scaffold attachment regions (MARs/SARs) DNA with an unmethylated 5’-CAC/GTG-3’ consensus sequence. In this study, we investigated how MeCP2 recognizes unmethylated 5’-CAC/GTG-3’ motif containing DNA by binding and structural studies. We found that MeCP2-MBD binds to MARs DNA with a comparable binding affinity to mCG DNA, and the MeCP2-CAC/GTG complex structure revealed that MeCP2 residues R111 and R133 form base-specific interactions with the GTG motif. For comparison, we also determined crystal structures of the MeCP2-MBD bound to mCG and mCAC/GTG DNA, respectively. Together, these crystal structures illustrate the adaptability of the MeCP2-MBD toward the GTG motif as well as the mCG DNA, and also provide structural basis of a biological role of MeCP2 as a transcription activator and its disease implications in Rett syndrome.

## Introduction

The discovery of MeCP2 as a nuclear, methyl-CpG DNA binding protein in 1992 is a major breakthrough in the field of DNA methylation^1^. In addition to its methyl-cytosine-binding domain (MBD)^2^, the MeCP2 protein also contains a NCoR/SMRT interaction domain (NID), which recruits the NCoR/SMRT corepressor complex to chromatin^3,4^, hence linking DNA methylation to histone deacetylation and transcriptional repression. The *MECP2* gene is located on the X chromosome in band 28 (Xq28), and ubiquitously expressed. In neurons, MeCP2 protein levels are comparable to histones^5^, suggesting a critical role in neuron development, and consistent with a link between *MECP2* mutations and Rett syndrome (RTT), a progressive X-linked neurological disorder almost exclusively found in girls^6^. Mutations or duplications of *MECP2* have also been found in patients with other neurological and autoimmune disorders^7^.

MeCP2 binding to chromatin was initially thought to depend on mCG density^5^. Subsequently, MeCP2 was also detected bound to non-CG methylated DNA^8–11^, and MeCP2 binding to both CG and non-CG methylated DNA contributes to transcriptional repression^8–10,12^. However, two other studies in either human neuronal SH-SY5Y cells^13^ or mouse hypothalamus^14^ reveal that the majority of genes are actively expressed with MeCP2 bound to their sparsely methylated promoters. Hence, MeCP2 may serve as a transcription repressor or activator in DNA methylation dependent or independent manners, respectively. This raises the question of how MeCP2 is able to bind to both methylated and unmethylated DNA. Interestingly, even before MeCP2 was identified as a methyl-CG binding protein, Attachment Region Binding Protein (ARBP), the chicken ortholog of human MeCP2 whose MBD domains share 100% sequence identity^15^, was found attached to Matrix/scaffold attachment regions (MARs/SARs) that lacked any methylated CG dinucleotides^15–17^. DNase I footprint analysis coupled with mutagenesis analysis showed that chicken MeCP2 specifically recognizes a sequence motif of 5’-GGTGT-3’ on MARs/SARs^15,17^. The central GTG trinucleotide is critical for the high affinity binding of the MARs DNA to chicken MeCP2, and mutating either of its guanine bases significantly reduced its binding to MeCP2^15,17^. Genes flanked by MARs exhibit elevated expression to some different extents^18–22^.

The complex structure of the MeCP2 MBD with symmetrically methylated CG DNA has been reported before^23^, however it is still unclear how MeCP2 recognizes unmethylated MARs DNA. In this study, we demonstrated the binding characteristics of MeCP2 to MARs DNA using ITC binding assays and structural analysis coupled with mutagenesis studies. Our findings presented here explain why MeCP2 selectively binds GTG-containing DNA as well as mCG DNA, and also provide a mechanistic insight into the biological role of MeCP2 as a transcription activator and its disease implications in Rett syndrome.

## Results

### MeCP2 binds unmethylated MARs DNA with high affinity

The selectivity of the MeCP2-MBD toward methylated CG DNA has been quantified using different synthetic DNA sequences and structural analysis of the MeCP2-MBD in complex with a mCG-containing DNA fragment from the promoter of the mouse brain-derived neurotrophic factor (*BDNF*)^23,24^. Nevertheless, the physiological context of the MeCP2-MBD binding to unmethylated DNA has not been settled. By means of electrophoretic mobility shift assay (EMSA), the Strätling group showed that ARBP (or chicken MeCP2) binds to a fragment of chicken lysozyme MARs by recognizing the DNA’s central 5’-GTG-3’ motif^15–17^. A similar ARBP-binding site has also been shown on mouse satellite DNA^15^. Other groups, however, propose that the binding of MeCP2 to unmethylated DNA might be incidental to the MBD’s core function of binding to methylated DNA^25^, i.e., the incidental binding of MeCP2 to unmethylated DNA would facilitate eventual binding to methylated DNA *in vivo*^1^.

In order to reconcile the contradictory interpretations of MeCP2 binding to unmethylated DNA, we first measured the affinity of the MeCP2-MBD to a 16mer (5’-GTGCAG<u>GTG</u>TCCTTAA-3’) DNA fragment from a chicken lysozyme MARs by ITC at approximately 2.8 µM (Fig. 1a), which is comparable to its affinity to an mCG DNA fragment (5’-GCCACTC<u>mCG</u>GAGTGGC-3’: ~1.8 µM)^26^. AT-rich sequences have often been found near the 5’-GTG-3’ consensus sequence and are thought to enhance their binding to MeCP2^15,16^. Interestingly, replacing the surrounding AT sequence of the CAC/GTG trinucleotide in the MARs with a sequence similar to that of the 16mer mCG DNA (5’-GCCCTG<u>GTG</u>TAGTGGC-3’: ~3.8 µM, Fig. 1a, b), barely changed its affinity, confirming that MeCP2 binds to the MARs DNA mainly through the central CAC/GTG motif^17^.

**Fig. 1.**
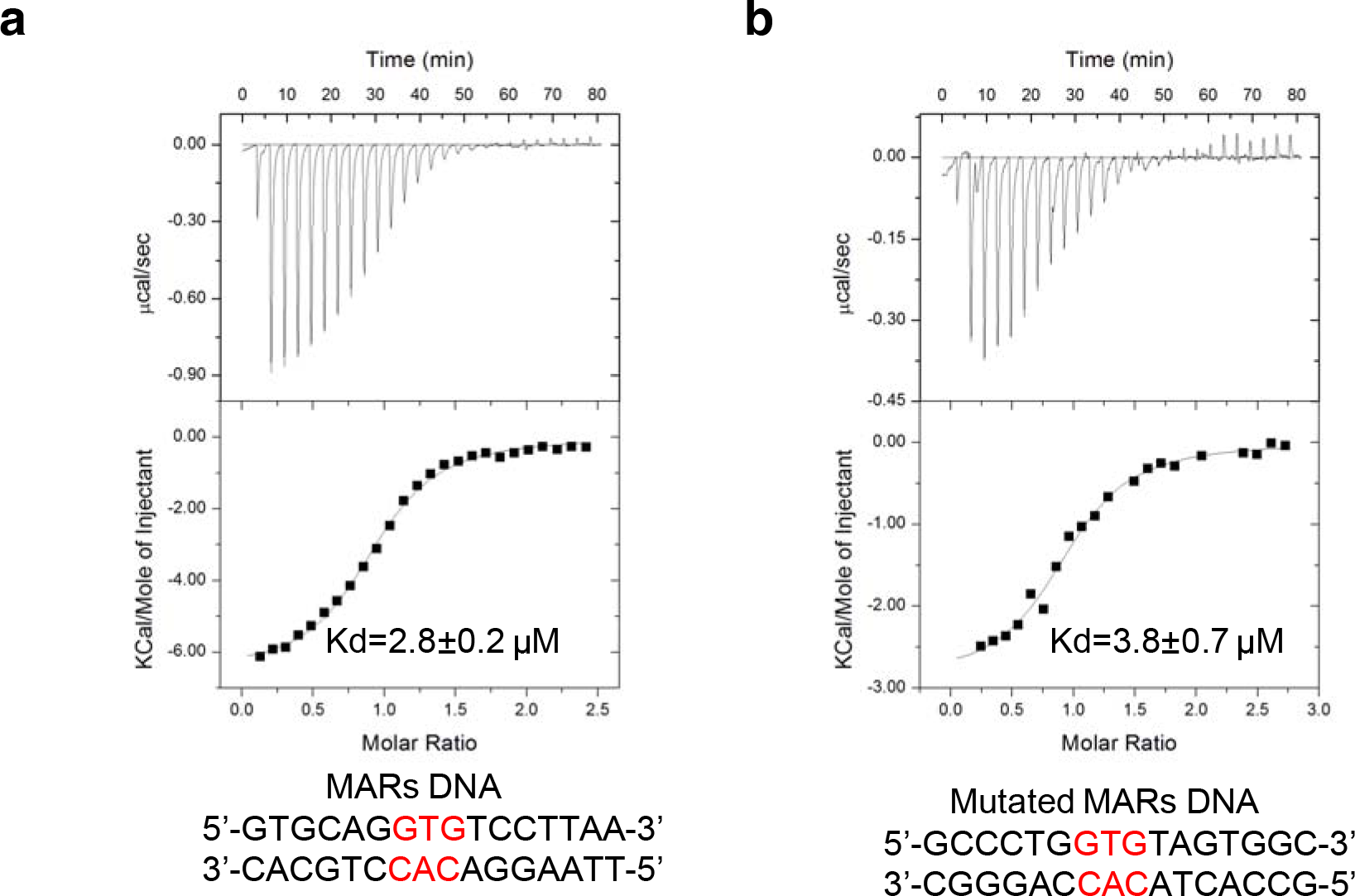
MeCP2-MBD specifically binds to unmethylated CAC/GTG motif containing DNA fragment from MARs. The ITC binding curves of the MeCP2-MBD to a native MARs DNA fragment (a) and a mutated MARs DNA fragment (b).

### Structural basis of MeCP2 binding to unmethylated CAC/GTG DNA

In order to understand the structural basis of the specific recognition of unmethylated DNA by the MeCP2-MBD, we attempted crystallization of the MeCP2-MBD in complex with different unmethylated DNA fragments that were centered a CAC/GTG trinucleotide motif, and obtained high resolution complex crystals with a 12mer CAC/GTG DNA. As a comparison, we also determined the complex structure of the MeCP2 MBD with a 12mer palindromic mCG DNA. Detailed data collection and structure refinement statistics are listed in Table 1.

**Table 1.** Data collection and refinement statistics of the MeCP2 MBD in complex with different DNA sequences

|  | MeCP2 (aa 80–164) | MeCP2 (aa 77–166) | MeCP2 (aa 77–167) |
| --- | --- | --- | --- |
| **DNA sequence** | 5'-GCCAC<u>mCG</u>GTGGC-3' | 5'- GCCTA<u>CAC</u>TCCG-3'<br>3'- CGGAT<u>GTG</u>AGGC-5' | 5'- GCCTA<u>mCAC</u>TCCG-3'<br>3'- CGGAT<u>GTG</u>AGGC-5' |
| **Crystallization Buffer** | 25% PEG3350, 0.1 M ammonium sulfate, 0.1 M tris, pH 8.5 | 30% PEG 2000 MME, 0.2M KBr | 0.1 M Imidazole and MES monohydrate (acid), pH 6.5, 0.018 M Magnesium chloride hexahydrate and 0.018M Calcium chloride dihydrate, 12.5% v/v MPD; 12.5% PEG 1000; 12.5 % w/v PEG 3350. |
| **Data Collection** |  |  |  |
| Space group | P1 | P2<sub>1</sub> | P2<sub>1</sub>2<sub>1</sub>2<sub>1</sub> |
| Cell dimensions |  |  |  |
| a,b,c [Å] | 40.75, 47.36, 54.62 | 29.72, 90.18, 40.52 | 40.47, 51.98, 66.30 |
| α, β, γ [°] | 68.15, 89.83, 65.98 | 90, 92.57, 90 | 90, 90, 90 |
| Resolution [Å] | 39.85-2.30 (2.38-2.30) | 45.09-1.80 (1.83-1.80) | 40.91-1.60 (1.63-1.60) |
| Completeness [%] | 96.3 (84.2) | 98.6 (94.1) | 99.8 (97.3) |
| Rsym | 0.068 (0.449) | 0.047 (1.111) | 0.060 (1.835) |
| I/sigmaI | 10.1 (2.4) | 12.0 (0.9) | 17.5 (0.7) |
| Redundancy | 2.2 (2.1) | 3.3 (3.0) | 6.1 (4.1) |
| **Refinement** |  |  |  |
| Resolution [Å] | 39.85-2.30 | 45.09-1.80 | 40.90-1.65 |
| Reflections used | 13799/785 | 18390/1109 | 15648/1773 |
| No. atoms/ B-factor [Å<sup>2</sup>] | 2069/68.0 | 1703/41.9 | 1157/32.3 |
| Protein | 1090/69.1 | 1169/43.9 | 585/29.3 |
| DNA | 976/66.9 | 485/37.5 | 498/36.4 |
| Water | not applicable | 41/38.1 | 56/27.7 |
| R work/free | 0.231/0.286 | 0.226/0.256 | 0.226 (0.255) |
| RMSD bonds [Å]/angles [°] | 0.016/1.7 | 0.011/1.5 | 0.011/1.7 |

\*Values in parentheses are for highest-resolution shell.

In the MeCP2-CAC complex structure (Fig. 2a-c), the MeCP2-MBD displays the canonical fold of a three-stranded β-sheet with a C-terminal α-helix packed against it. The β-sheet inserts into the major groove of the CAC DNA, consistent with the MeCP2-mCG structure presented here and other published MeCP2-mCG structures (Fig. 2d-f)^23,27^. In the MeCP2-CAC crystal structure, bases of only the CAC/GTG motif directly interact with MeCP2, confirming the importance of the central CAC/GTG trinucleotide in binding to MeCP2 (Fig. 2a-c). Arginine finger residues 111 and 133, which interact with the critical CAC/GTG motif all reside in the β-sheet core (Fig. 2a-c). Specifically, R111, rigidified by a salt bridge with D121, recognizes the TG dinucleotide of the G<u>TG</u> motif in a stair motif binding mode (Fig. 2a-c). R133 forms hydrogen bonds with the bases of the <u>GT</u>G motif’s 5’-teminal GT dinucleotide (Fig. 2a-c). In the MeCP2-mCG structure, on the other hand, arginine fingers R111 and R133 each recognize one mCG dinucleotide from the complementary DNA strands to form a pseudo-symmetric protein-DNA interface (Fig. 2d-f)^23^.

**Fig. 2.**
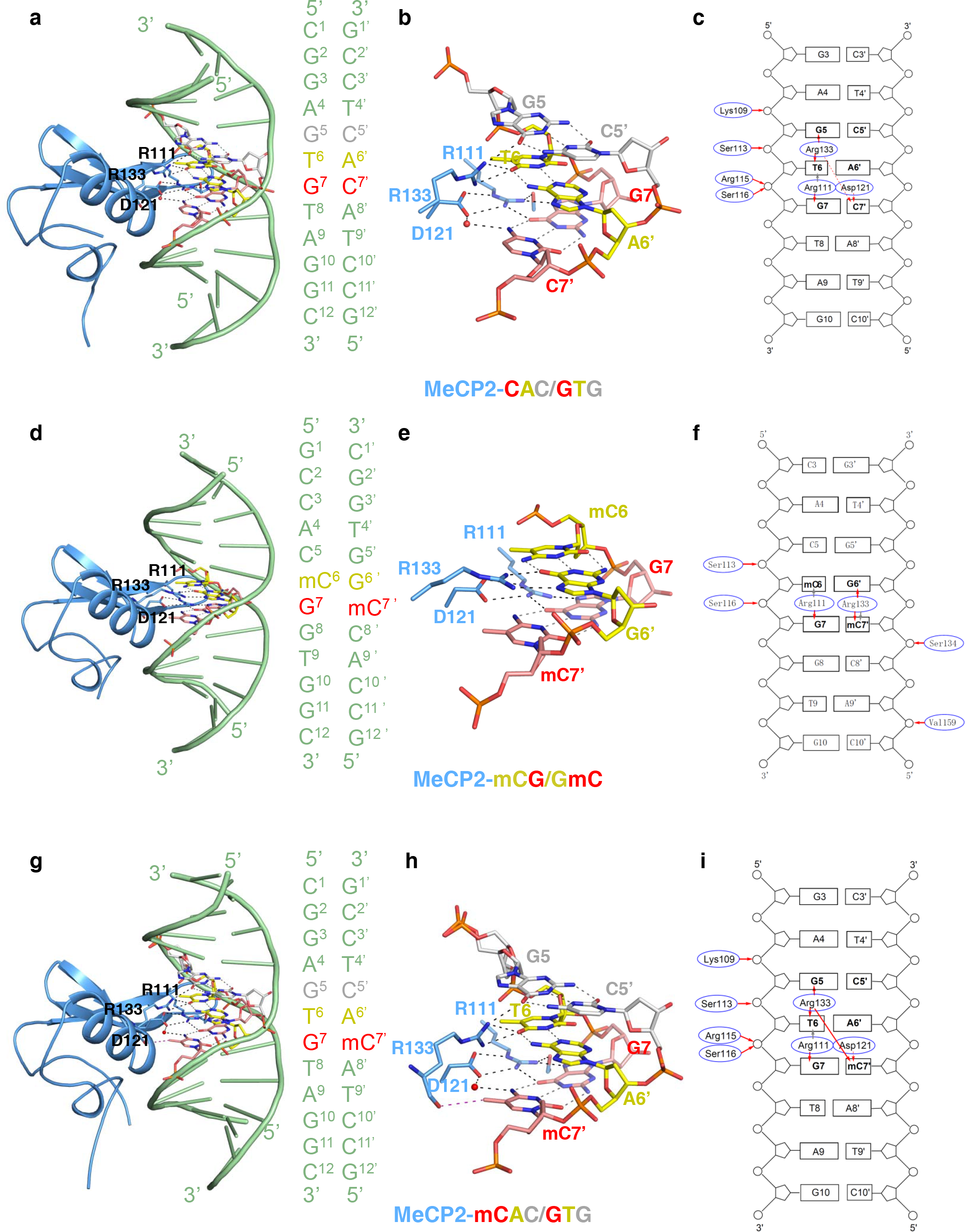
Structural basis for the recognition of CAC, mCG and mCAC by MeCP2-MBD. **(a, d and g)** Overall structures of MeCP2-MBD in complex respectively with CAC, mCG and mCAC DNA, respectively. The protein part is shown in blue cartoon representation, while the DNA ligand is shown in green cartoon representation. The nucleotides involved in base interactions are shown in stick models: G5-C5’ (grey), mC6-G6’ or TC6-A6’ (yellow), G7-mC7’ or G7-C7’ (red). The base interacting protein residues in MeCP2 are shown in stick models and water molecules are shown in red spheres. The hydrogen bonds formed between protein residues and DNA are marked as black dashed lines, while the grey dashed lines represent the hydrogen bonds between base pairs. In **(g)**, the non-conventional C-O hydrogen bond between 5-methyl group of the methylated cytosine in the mCAC/GTG sequence and the main chain carbonyl oxygen of R133 was colored as purple. **(b, e and h)** Detailed interactions of different DNA with MeCP2-MBD. The interacting residues and DNA bases are shown in the same mode as in (a, d and g), respectively. The hydrogen bonds formed by protein residues and DNA are marked as black dashed lines, while the grey dashed lines represent the hydrogen bonds between base pairs. In (h), the non-conventional C-O hydrogen bond between 5-methyl group of the methylated cytosine in the mCAC/GTG sequence and the main chain carbonyl oxygen of R133 was colored as purple. **(c, f and i)** Schematic diagram of the detailed interactions between MeCP2-MBD and different DNA. Direct and water-mediated hydrogen bonds are indicated by solid and dash red arrows, respectively. The stacking interactions between Arginine fingers and bases are indicated by grey arrows.

### Structural basis of a mild preference of MeCP2 toward methylated mCAC over CAC DNA

CG sites are the major cytosine methylation sites, but non-CG (CH, H=A, T, or C) sites are also subject to cytosine methylation, especially in embryonic stem cells and neurons^8,28^. Among the non-CG sites, the trinucleotide CAC motif is the most prominent modification site, forming mCAC^8^. It was reported recently that MeCP2 binding to chromatin is dependent on the total methylation density of both mCG and mCAC in mouse brain^10^. Our ITC results indicate that the MeCP2-MBD binds to a mCAC DNA with an affinity of ~0.9 µM, slightly tighter than to the MARs DNA (Fig. 1a, b)^26^. In order to address the recognition difference of mCAC and CAC by MeCP2, we determined a complex structure of the MeCP2-MBD with a 12mer mCAC/GTG DNA (Table 1).

The MeCP2-mCAC/GTG structure is very similar to that of MeCP2-CAC/GTG, with an RMSD of 0.4 Å over all the aligned Cα atoms (Fig. 2g-i and 3a) and largely conserved GTG recognition (Fig. 2h, i). However, the mCAC 5-methyl group points toward the main chain carbonyl oxygen of R133 with a C-O distance of approximately 3.4 Å (Fig. 2g, h and 3a). This arrangement may represent a weak hydrogen bond, similar to those involving side chain methyl groups in proteins^29^, and might explain why MeCP2 binds to mCAC slightly more tightly than to CAC DNA (Fig. 1a, b)^26^. In both MeCP2-mCAC/CAC structures, the GTG trinucleotide appears to be the dominant driver of protein-DNA binding (Fig. 2b, h), which is consistent with the observation that 5-hydroxymethylation of a central hmCA motif has little or no effect on recognition by MeCP2^11^.

**Fig. 3.**
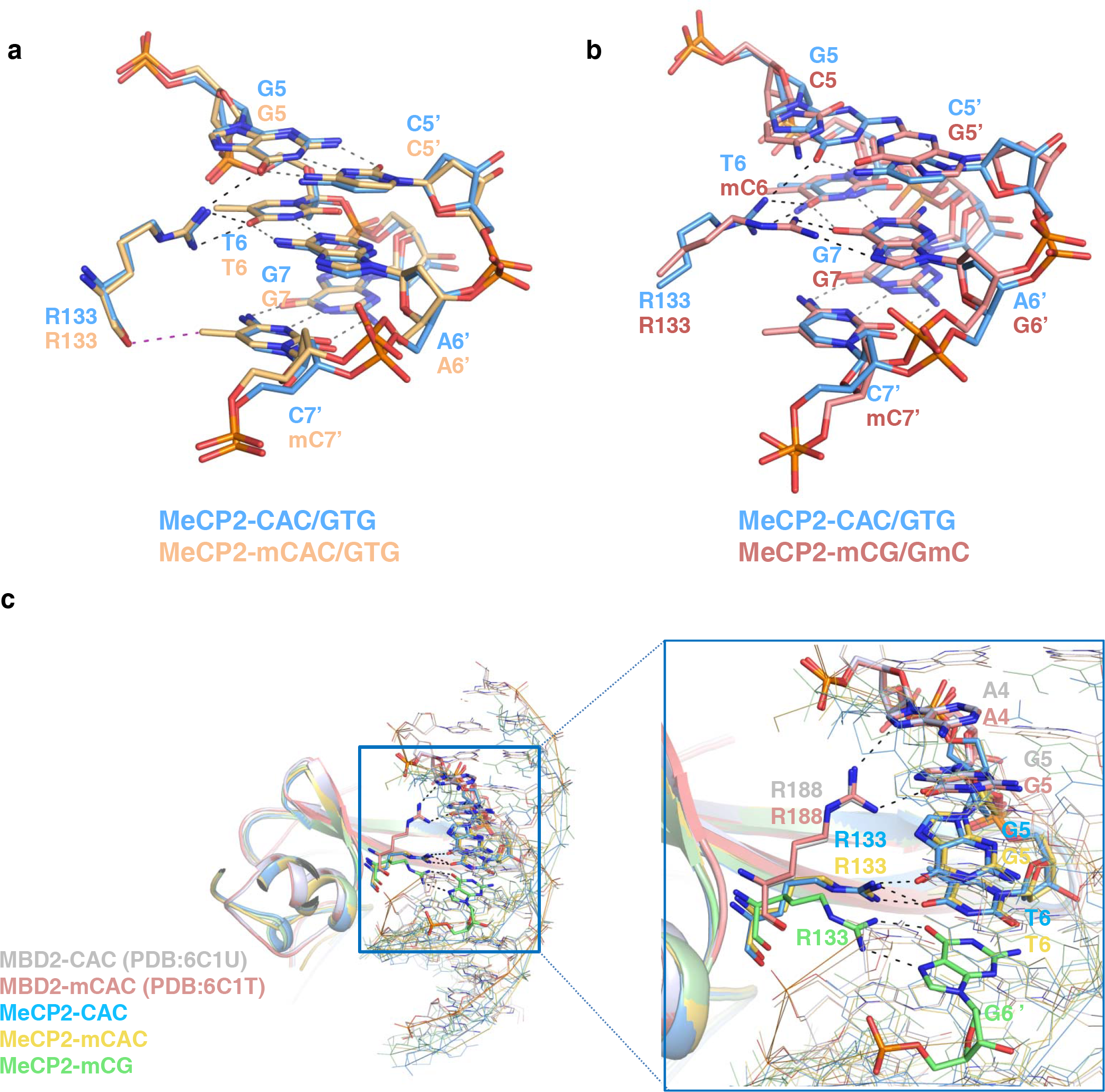
Structural comparisons of MeCP2 and MBD2 MBDs in complex with different DNA. **(a)** Structural comparisons of MeCP2-MBD in complex with CAC/GTG and mCAC/GTG DNA. Hydrogen bonds formed by protein and DNA are shown as the black dashed lines, the interaction between DNA base pairs are shown as grey dashed lines. The non-conventional C-O hydrogen bond between 5-methyl group of the methylated cytosine in the mCAC/GTG sequence and the main chain carbonyl oxygen of R133 was colored as purple. **(b)** Structural comparisons of MeCP2-MBD in complex with CAC/GTG and mCG DNA. Hydrogen bonds formed by protein and DNA are shown as the black dashed lines, the interaction between DNA base pairs are shown as grey dashed lines. **(c)** Structural comparisons of MBD2-CAC (grey), MBD2-mCAC (pink), MeCP2-CAC (blue), MeCP2-mCAC (Yellow), and MeCP2-mCG (green) complexes. Arg133 of MeCP2 and Arg188 of MBD2 are shown in sticks. Hydrogen bonds formed by protein and DNA are shown as the black dashed lines.

Recently we also solved the MBD2-mCAC/GTG and MBD2-CAC/GTG structures^26^. The major difference between the MeCP2 and MBD2 structures is that the second arginine finger R188 in MBD2 (corresponding to R133 in MeCP2) does not interact with the thymine base of the GTG motif. Instead, it forms hydrogen bonds with the first guanine of the GTG motif and the preceding adenine (Fig. 3c)^26^. In the MeCP2-mCAC/CAC structures, the second arginine finger R133 closely interacts with the GT dinucleotide of the GTG motif (Fig. 3c), which is consistent with slightly stronger binding of GTG-containing DNA to MeCP2 than to MBD2^26^. These two recognition modes of the second arginine finger are diverse from that observed in palindromic mCG complexes, in which the second arginine finger contributes to a pseudo-symmetric protein-DNA interface, as described above and also observed in other MBD-mCG structures (Fig. 3b, c)^23,27,30–33^. These observations confirm our previous assertion that the second arginine finger, such as R133 in MeCP2, is more flexible than the first arginine finger, like R111 in MeCP2, to enable a broader binding DNA selectivity spectrum^26^.

### Effects of Rett Syndrome mutations of MeCP2 on its binding to mCG and GTG DNA

Based on our structural studies, R111 and R133 of MeCP2 are the two most critical residues in DNA binding. Mutations of R111 and R133 have been found in Rett Syndrome patients, and R133 is a Rett Syndrome hotspot mutation point^34^. It has been reported that both R111G and R133C mutations of MeCP2 weaken its binding to mCG DNA as measured by gel mobility shift assays^35^. We investigated here how the R111 and R133 mutations of MeCP2 affect its binding to both mCG and GTG DNA by measuring the binding affinities of the R111G, R111A, R133C and R133A mutants to various DNAs. As expected, the R111G and R111A mutations of MeCP2 disrupted its binding to either GTG or methylated DNA (Fig. 4a, b), consistent with our previous report that the first arginine finger is the major contributor to DNA binding by MBDs^26^. Neither the R133C nor R133A mutants of MeCP2 measurably bound to CAC/GTG DNA, but they still retained significant binding to methylated DNAs, including mCAC/GTG and mCG DNA (Fig. 4c, d). As mentioned above, the side chain of R133 forms hydrogen bonds with the GT dinucleotide in the GTG motif (Fig. 2b, h). However, in the MeCP2-mCAC/GTG structure, the main chain of R133 may also form an additional, weak hydrogen bond with the methyl group from the methylated cytosine (Fig. 2h and 3a), which would be absent in the MeCP2-CAC/GTG structure (Fig. 2b and 3a). Although mutating R133 into cystine or alanine disrupts its side chain interactions, interactions of the main chain, such as one of the carbonyl oxygen, would not be directly affected. Thus, it seems that the concurrent loss of both side and main chain interactions may render R133 mutants incapable of binding unmethylated CAC/GTG DNA. One potential mechanism of the MeCP2 mutations leading to the Rett Syndrome may be due to their loss of DNA binding ability. The observations that the R133 mutants are still able to bind to mCG and mCAC/GTG, but not to CAC/GTG, led us to speculate that the unmethylated CAC/GTG DNAs are a physiological ligand of MeCP2 in neurons, and the disrupted binding between them might play a role in the Rett Syndrome. This implication needs further investigation in the future.

**Fig. 4.**
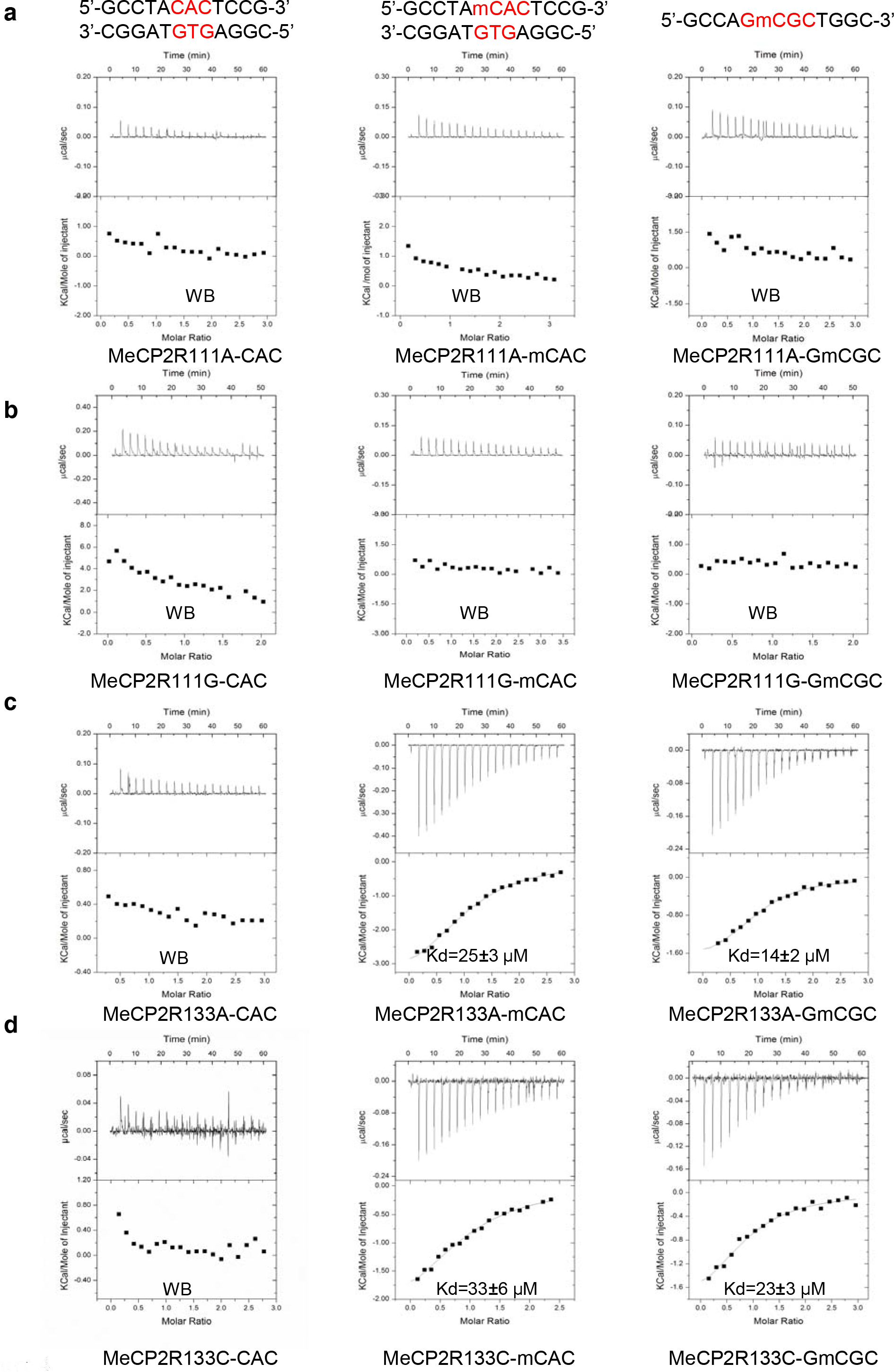
The R111A, R11G, R133A and R133C mutations of MeCP2 reduce its DNA binding ability. **(a)** ITC binding curves of the MeCP2 R111A mutant to different DNA. **(b)** ITC binding curves of the MeCP2 R111G mutant to different DNA. **(c)** ITC binding curves of the MeCP2 R133A mutant to different DNA. **(d)** ITC binding curves of the MeCP2 R133C mutant to different DNA. WB: Weak binding.

In addition to R111 and R133, many other RTT-associated mutations have been mapped to the MeCP2-MBD (Fig. 5a)^35–38^, and most of them have been reported to reduce the MeCP2 binding to methylated DNA^35^. Based on the MeCP2-DNA complex structures, RTT-associated mutations in MBD can be classified into three groups: 1) are directly involved in base-specific interactions, such as the arginine fingers R111 and R133; 2) contact the DNA backbone by providing hydrogen bonding, electrostatic and/or Van Der Waals Interactions, such as S134C, K135E and T158M; and 3) not directly interact with the DNA ligand, and may affect binding indirectly by changing the overall protein stability or solubility, such as R106W, Y141, and D156^35,37^. To test how these mutations affect their binding to the CAC DNA, we chose 6 high-frequency mutation points (R106W, S134C, K135E, Y141C, D156E and T158W), and carried out ITC assays. Of the three DNA-proximal mutants, the S134C and T158M mutants exhibited significantly reduced or disrupted binding, but the K135E only had slightly reduced binding (Fig. 5b). Although K135 is near DNA, but its side chain is exposed to the solvent, and its ε-ammonium group is not in contact with the DNA, and is disordered in our structures (Fig. 5a). Mutating R106W, Y141C, or D156E caused loss of the DNA binding ability (Fig. 5b), potentially due to reduced protein stability because all of these residues form extensive interactions with surrounding residues. Therefore, most of the RTT-associated mutations in the MeCP2-MBD cause Rett syndrome by diminishing or disrupting the MeCP2’s ability in binding to its DNA ligands.

**Fig. 5.**
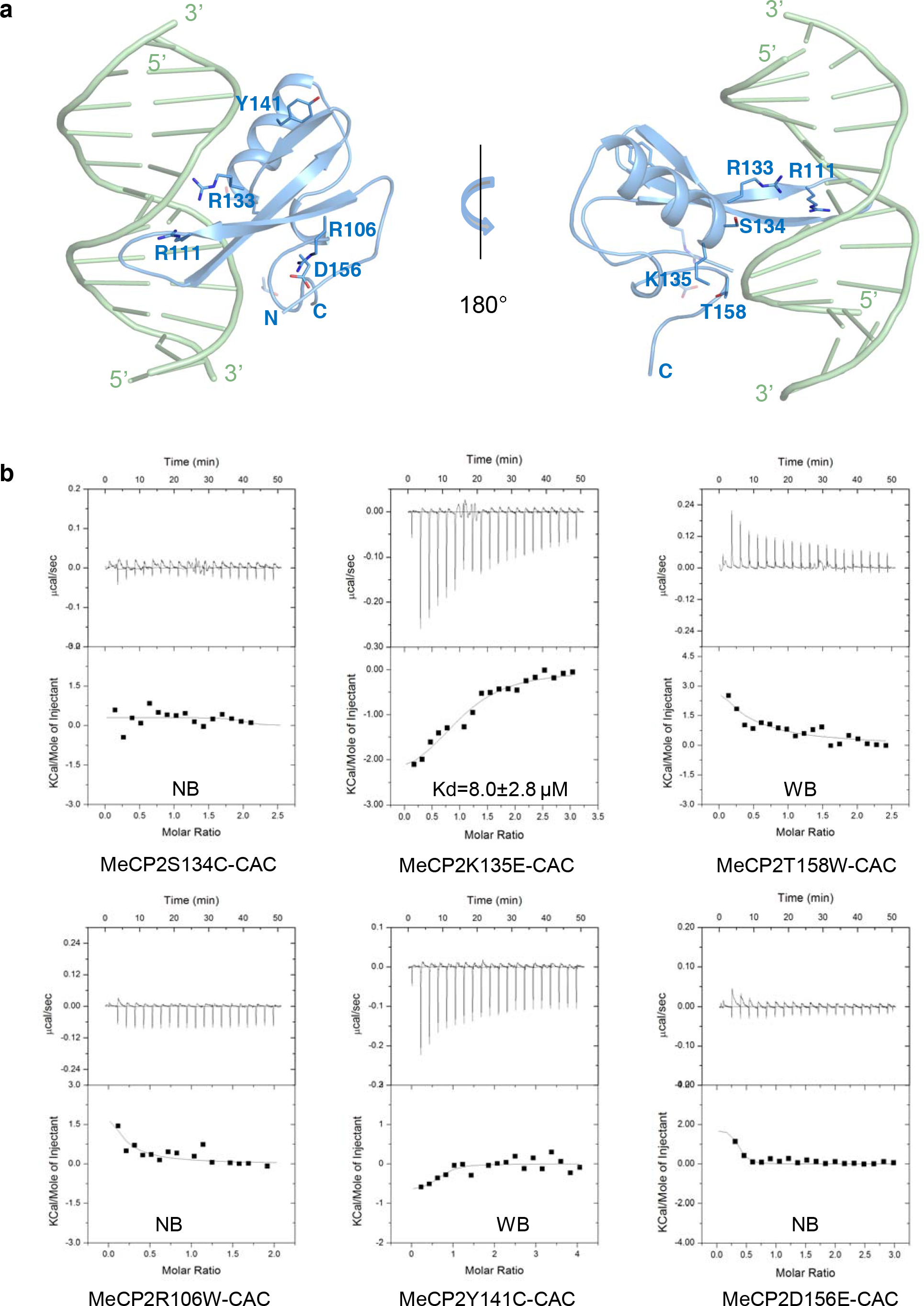
Effect of Rett Syndrome mutations of MeCP2-MBD on its binding to CAC/GTG DNA. **(a)** Locations of some high-frequency mutations on the MeCP2-MBD. The protein (blue) and CAC DNA (green) are shown in cartoon representation, and mutated residues in the MeCP2-MBD are shown as stick models. **(b)** ITC binding curves of 6 chosed MeCP2-MBD mutations to CAC DNA. WB: Weak binding. NB: No detectable binding.

## Discussion

As MeCP2 recruits histone deacetylases (HDAC)^4^ and histone methyltransferases^39^ to methylated chromatin loci, a role for MeCP2 in transcription repression has been proposed^4,39^. A recent ChiP-seq analysis has revealed that the genome wide distribution of MeCP2 is determined by the percentage of GC, not necessarily by the density of methylated CG dinucleotide^40^, suggesting that the location of MeCP2 is not always correlation with heavily methylated CG. Methylated non-CG motifs, such as mCAC, were found to bind MeCP2 in brain^9,10^. MeCP2 also associates with unmethylated CAC/GTG containing DNA fragments, including the MARs/SARs DNA and mouse satellite DNA *in vitro*^15–17^. MARs/SARs DNA elements, which are evolutionarily conserved from yeast to human, might serve as chromatin loop anchors and play a role in the regulation of gene expression^16,41,42^. In interphase MCF7 cells, MeCP2 was found to be associated with four clones derived from putative MARs, but just one clone from CG islands^43^. MeCP2 was found to activate the majority of its target genes in mouse hypothalamus, and corresponding promoters were not necessarily rich in mCG sites^14^. Clearly, MeCP2 association requires neither the CG motif nor cytosine methylation, significantly widening the repertoire of potential ligands beyond initial predictions.

Our ITC data confirmed that MeCP2 binds to DNA that contains the CAC/GTG trinucleotide motif, which was originally identified in some MARs/SARs DNA as the ligand of the MeCP2 chicken ortholog^16^. Our crystal structures revealed that bases of only the GTG trinucleotide interact with the residues R111 and R133 in MeCP2 MBD and that cytosine methylation on the complementary strand is not essential. Additionally, cytosine methylation in mCAC introduces an additional contact with a potential hydrogen bond acceptor on MeCP2 R133, the second-most commonly mutated MBD residue in RTT patients. The role of R133 in base recognition and the possibility of a weak interaction formed by the MBD main chain and a putative cytosine 5-methyl group together would explain our failure to detect binding between MeCP2 R133 mutants and unmethylated GTG DNA, while we still detected binding for mCAC and mCG DNA (Fig. 4c, d). The observations that 1) the R133 mutants are still able to bind to mCAC and mCG DNA but lack binding to CAC DNA, and 2) most of MeCP2-controlled genes are sparsely methylated lead us to speculate that the unmethylated CAC/GTG motif containing DNA is one major physiological ligand of MeCP2, which need to be further evaluated in the future.

## Methods

### Protein expression and purification

Human MeCP2 (aa. 80–164) was subcloned into pET28-MHL expression vector to generate N-terminal His tag fused protein. The MeCP2-MBD mutants were obtained by Quick Change site-directed mutagenesis (Agilent Technologies) using the MeCP2 (aa. 80–164)-pET28-MHL plasmid as template. For crystallization, two other MeCP2-MBD (aa. 77-167 and aa. 77-166) fragments were subcloned into the pNIC-CH expression vector, respectively.

The recombinant proteins were overexpressed in *Escherichia coli* BL21 (DE3)-RIL strains and induced with 0.35 mM isopropyl-β-d-thiogalactopyranoside (IPTG) at 16°C for overnight. The cell pellet was collected, and further dissolved in a buffer containing 20 mM Tris-HCl, pH 7.5, 500 mM NaCl, 0.5 mM PMSF and 5% glycerol. The lysed cell supernatants were incubated with Ni-NTA resin (Qiagen) for 30 min before washing with lysis buffer with an additional 10 mM imidazole. Finally, the purified proteins were eluted using 20 mM Tris-HCl, pH 7.5, 500 mM NaCl and 300 mM imidazole. For crystallization, the His tag in MeCP2 (aa. 80–164) fusion proteins was removed by tobacco etch virus (TEV). The proteins were further purified by anion-exchange column and gel filtration column (GE Healthcare). Finally, the purified proteins for ITC experiments were concentrated to 10 mg/ml in 20 mM Tris-HCl, pH 7.5 and 150 mM NaCl. For crystallization, 1 mM DTT was added in the ITC buffer.

### ITC Binding assay

All the DNA oligos used for ITC and crystallization experiments were synthesized by IDT (Integrated DNA Technologies, USA) and processed as described before^26^. Simply, the single DNAs were dissolved in the same buffer as the target proteins, containing 20 mM Tris-HCl, pH 7.5 and 150 mM NaCl, whose final pH was adjusted to around 7.5 using NaOH. Then the single strand DNA was annealed into DNA duplex^44^. The concentrations of MeCP2 and DNA oligos used in ITC experiment are ranging from 30 to 80 µM and 0.5 mM to 1 mM, respectively. The concentrations of proteins and DNA are the average concentrations that were determined by Nano Drop (Thermo Scientific) over triple measurements. ITC measurements were carried out using MicroCal ITC or ITC200 (GE Healthcare) at 25 °C. The data were fitted using Origin 7.0 (MicroCal Inc.) with one-site binding model of the ITC data analysis module. The standard error of each Kd value reflects the fitting error for the best ITC titration curve. The detailed thermodynamic parameters have been reported in the Supplemental Table 1.

### Crystallization

The purified proteins were mixed with DNA at a molar ratio approximately 1:1.3 by directly adding the DNA into the protein solution. After incubation on ice for 30 minutes, the protein–DNA reaction mixtures were crystallized using the sitting drop vapor diffusion at 18 °C. The detailed crystallization conditions for each MeCP2 MBD-DNA complex were summarized in Table 1.

### Data collection and structure determination

Crystals were soaked in a mixture of 1 part glycerol to between 4 and 6 parts of the relevant crystallization reagent prior to rapid immersion in liquid nitrogen for storage. Diffraction data were collected under cooling at the Advanced Photon Source (mCG complex), Canadian Light Source (CAC)^45^ or Advanced Light Source (mCAC). Structures were solved by molecular replacement with the program PHASER^46^. For the mCG complex, search models were derived from PDB entry 3c2i^23^ and a preliminary version of coordinates from PDB entry 6cnq^26^. Search models for the other complexes were derived from the mCG complex model. ARP/wARP^47^ was used to improve electron density maps for the mCG and mCAC complexes. Data collected on a rotating anode source from an additional, nearly isomorphous crystal were used to solve the structure of the mCAC complex, and DIMPLE (http://ccp4.github.io/dimple) scripts were used during refinement of that complex’s atomic model. Models were interactively rebuilt with COOT^48^ and refined with REFMAC^49^ or PHENIX^50^. Model geometry was evaluated with MOLPROBITY^51^. Data collection and refinement statistics were summarized in Table 1.

## Acknowledgements

GM/CA@APS has been funded in whole or in part by the National Cancer Institute (ACB-12002) and the National Institute of General Medical Sciences (AGM-12006). This research used resources of the Advanced Photon Source, a U.S. Department of Energy (DOE) Office of Science User Facility operated for the DOE Office of Science by Argonne National Laboratory under Contract No. DE-AC02-06CH11357. Research described in this paper was performed using beamline 08ID-1 at the Canadian Light Source, which is supported by the Canada Foundation for Innovation, Natural Sciences and Engineering Research Council of Canada, the University of Saskatchewan, the Government of Saskatchewan, Western Economic Diversification Canada, the National Research Council Canada, and the Canadian Institutes of Health Research. The Berkeley Center for Structural Biology is supported in part by the Howard Hughes Medical Institute. The Advanced Light Source is a Department of Energy Office of Science User Facility under Contract No. DE-AC02-05CH11231. The Pilatus detector on 5.0.1, was funded under NIH grant S10OD021832. The ALS-ENABLE beamlines are supported in part by the National Institutes of Health, National Institute of General Medical Sciences, grant P30 GM124169. The SGC is a registered charity (number 1097737) that receives funds from AbbVie, Bayer Pharma AG, Boehringer Ingelheim, Canada Foundation for Innovation, Eshelman Institute for Innovation, Genome Canada through Ontario Genomics Institute [OGI-055], Innovative Medicines Initiative (EU/EFPIA) [ULTRA-DD grant no. 115766], Janssen, Merck KGaA, Darmstadt, Germany, MSD, Novartis Pharma AG, Ontario Ministry of Research, Innovation and Science (MRIS), Pfizer, São Paulo Research Foundation-FAPESP, Takeda, and Wellcome. We also thank Dr. Zhang Delin at Center for Protein Research (CPR), Huazhong Agricultural University for technical support.

## AUTHOR CONTRIBUTIONS

K.L and J.M. conceived the project. M.L. and K.L. designed and performed crystallization and binding experiments and data analysis. W.T. determined the crystal structures. K.L and J.M. wrote the manuscript with significant contributions from the other authors.

